# Diazoxide/dibenozylmethane treatment mitigates spatial memory deficits and pathology and upregulates protective genes in an Alzheimer’s transgenic rat model

**DOI:** 10.1101/2023.09.01.555966

**Authors:** Charles Wallace, Giovanni Oliveros, Lei Xie, Peter Serrano, Patricia Rockwell, Maria E. Figueiredo-Pereira

## Abstract

**INTRODUCTION:** Alzheimer’s disease (AD) is a multifactorial disease for which therapeutic efficacy should benefit from a multi-target approach. Thus, we evaluated a combined chronic treatment with diazoxide (DZ) and dibenzoylmethane (DIB). DZ is a potassium channel activator. DIB counteract eIF2α-P-driven stress responses. The individual therapeutic benefits of each drug on attenuating neurodegeneration and apoptosis were previously examined, but not as a combined treatment.

**METHODS:** We evaluated the efficacy of chronic DZ/DIB treatment on TgF344-AD rats (Tg-AD) at 4- and 11-months of age and wild-type littermates. Spatial working memory was assessed with the radial 8-arm maze, and AD pathology by immunohistochemistry. We used RNA sequencing for transcriptome analysis.

**RESULTS:** DZ/DIB-treatment mitigated the spatial memory deficits as well as the buildup of hippocampal Aβ plaques and tau PHF exhibited by 11-month Tg-AD rats. The DZ/DIB-treatment had no effect on wild-type littermates. We did not detect AD-related deficits in untreated 4-month Tg-AD rats, but DZ/DIB-treatment altered their transcriptome. DZ/DIB-treatment of 4-month Tg-AD rats upregulated several genes normally downregulated in AD and/or aging. Expression of two potential early biomarkers for AD, *EGR2* (early growth response 2) and *HISIT1H2AA* (histone H2AA), were also altered by the DZ/DIB treatment in 4-month Tg-AD rats. The treatment reduced levels of eIF2α, a protein involved in abnormal translational repression and a contributing factor to neuronal loss.

**DISCUSSION:** This preclinical study represents the first report on the combined DZ/DIB-treatment. Besides the benefits of this treatment on spatial memory and AD pathology, we identified two potential early AD biomarkers. Furthermore, the DZ/DIB-treatment prevented downregulation of genes associated with AD and/or aging. Overall, our results strongly support that the combination DZ/DIB-treatment mitigates AD pathology. Evaluations across multiple AD-related models are warranted to further corroborate that the DZ/DIB combination is a candidate treatment for AD.

**Highlights:**

- DZ/DIB treatment improves cognitive deficits and AD pathology in TgF344-AD rats
- *EGR2* and *HISIT1H2AA* genes are potential early AD biomarkers in TgF344-AD rats
- DZ/DIB treatment of 4-month TgF344-AD rats blocks downregulation of AD/aging genes

**RESEARCH IN CONTEXT:** 1. Systematic Review: We reviewed the literature using traditional sources (e.g., PubMed) to assess the status of DZ and DIB treatment individually in preclinical studies in animal models. To our knowledge there is no data with this chronic combined treatment.

2. Interpretation: Treatment takes advantage of a polypharmacological approach to AD therapeutics, considering the multifactorial nature of AD. DZ and DIB individually have been successful in mitigating hippocampal AD pathology but their combined effects were unknown, particularly in a rodent model that shows a more complete disease progression that incorporates aging as a risk factor. The effects of the treatment go beyond previous data in individual DZ and DIB treatment, supporting the need for combination drug treatments to treat a multifactorial disorder.

3. Future Directions: Investigate the potential of DZ/DIB treatment to stop the progression or ameliorate AD pathology when administered at a later age, when pathology is first detected.

**Graphical Abstract:** 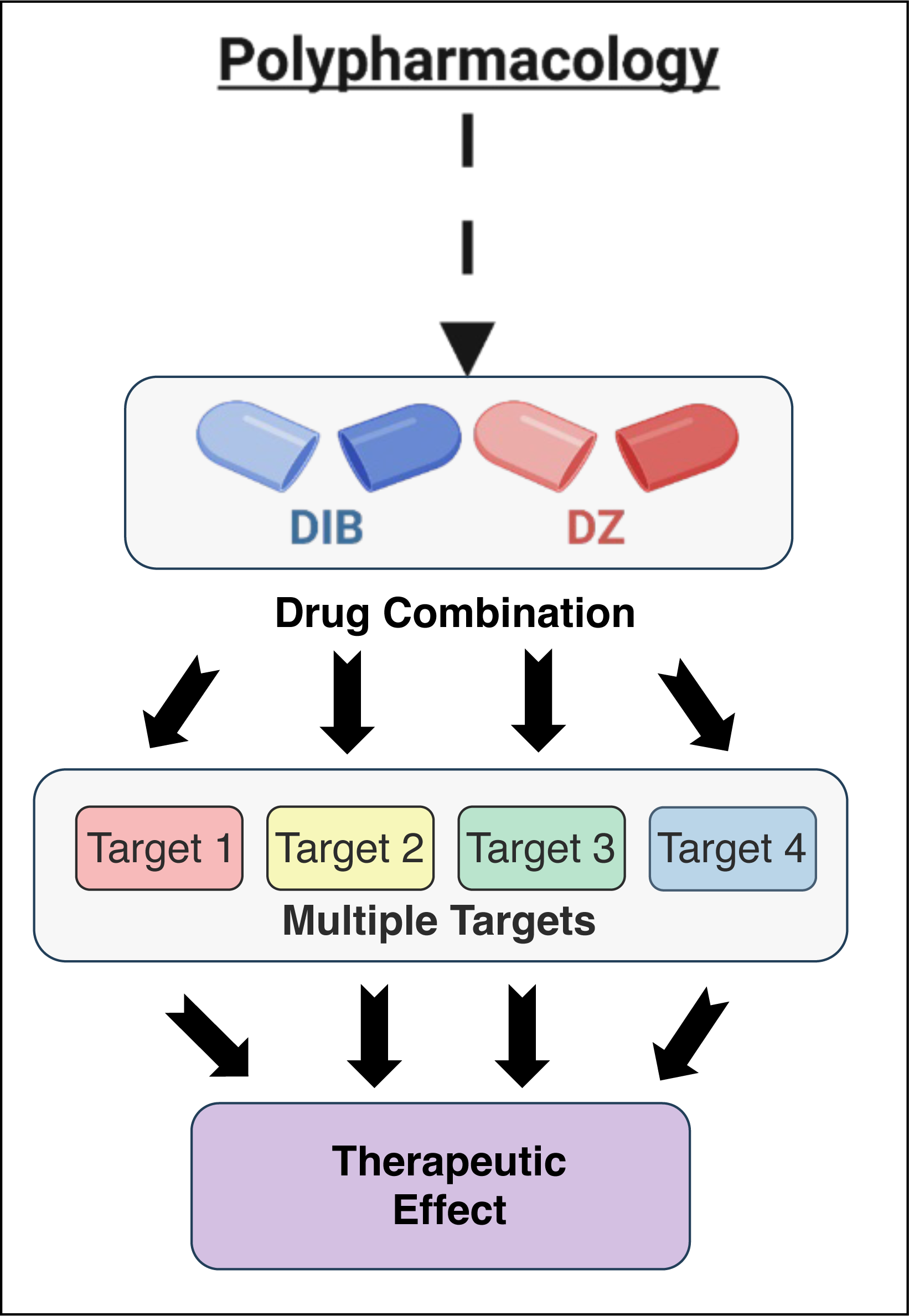

## 1. INTRODUCTION

Alzheimer’s Disease (AD) is the most common form of dementia^1^ and is a multifaceted neurodegenerative disease with aging being the major risk factor^2,3^. Mono-target therapeutics directed at amyloid plaques or tau tangles are common to treat AD^4^. Currently, while only seven FDA approved drugs have been approved as AD therapeutics, five of them have no effect on biological changes associated with AD, with the two most recent approved drugs (aducanumab and lecanemab) not recommended for all individuals living with AD due to serious side effects associated with treatment and limited safety data. Alternatively, the use of multi-target drugs that work on multiple biological pathways, often referred to as polypharmacology, is on the rise^5,6^. Through computational studies, drugs intended for other diseases can be predicted to be repurposed for AD^7^. Some benefits of drug repurposing include low development costs, known safety profiles, and bringing existing therapeutics to new patient populations^8^.

In this context, we investigated the effects of chronic treatment with diazoxide (DZ) and dibenzoylmethane (DIB) on the transgenic TgF344-AD (Tg-AD) rat model of AD^9^. DZ has been used for decades for its cardioprotective effects^10^. DZ is a benzothiadiazine derivative that is an agonist to mitochondrial ATP sensitive potassium channels, thus increasing intracellular K^+^ levels. and reducing abnormally elevated Ca^2+^ concentrations, which are commonly found in AD^11^. DZ alone prevented/mitigated cognitive deficits and histopathology in the 3xTg-AD mouse model of AD^12^, caspase-dependent apoptosis via BCL-2 activation and BAX inhibition in rat hippocampus^13^, and apoptosis in PC12 cells^14^.

DIB is a β-ketone analog of curcumin found in licorice among other plants^15^. DIB reverses eIF2α-P-mediated translational attenuation occurring in the unfolded protein response, which plays a critical role in controlling protein synthesis rates in cells^16^. This pathway is overactive in brains of AD patients, thus mitigating its activity with DIB could be beneficial^16^. DIB and its derivatives display beneficial effects, such as chemo-preventive, anti-cancer, anti-mutagenic, anti-inflammatory, liver protection, cellular stress preventive, iron-chelating abilities and is even protective against UV rays^15-23^. DIB also prevented hippocampal neuronal loss and improved memory deficits in transgenic prion-disease and tauopathy-frontotemporal dementia mouse models^16^. Interestingly, DIB was explored in models of cancer with various combination drug approaches^21,22^.

We investigated the therapeutic potential of a DZ/DIB combination in an AD context because each of these drugs used separately prevented the progression of neurogeneration in preclinical rodent models^12,16^. We evaluated the DZ/DIB treatment with TgF344-AD (Tg-AD) rats and their wild-type (WT) littermates. Tg-AD rats develop a wide array of AD pathologies, including cognitive deficits, Aβ plaques, neurofibrillary tangles (NFTs), neuroinflammation and neuronal loss, in an age-dependent progressive manner^9^.

Overall, our studies demonstrate that the DZ/DIB treatment mitigates spacial memory deficits and hippocampal Aβ-plaque and neurofibrillary tau-tangle burden in Tg-treated compared to untreated rats. Moreover, at 4 months of age (pre-pathology stage), the expression levels of the *EGR2* and histone *H1H2AA* genes were altered in female Tg-AD rats, suggesting that they could be early AD biomarkers. Our results strongly support that the DZ/DIB treatment is an effective strategy to mitigate AD pathology due to its multi-target approach that affects multiple signaling pathways.

## 2. MATERIAL AND METHODS

### 2.1 TgF-344 AD rats

Fisher transgenic 344-AD (Tg-AD) rats expressing human Swedish amyloid precursor protein (APPsw) and Δ exon 9 presenilin-1 (PS1-ΔE9) both driven by the prion promoter^9^, were purchased from the Rat Resource and Research Center (RRRC, Columbia, MO). Tg-AD (n = 55, Table 1) and WT (n = 50, Table 1) rats of both sexes were housed in pairs upon arrival and maintained on a 12-hour light/dark cycle with food and water available *ad libitum* (Figure 1). All animal procedures were approved by the IACUC at Hunter College. For rigor and reproducibility in accordance with the ARRIVE guidelines, all researchers were blinded to genotype and drug dose during study execution and through data analysis^24^.

**Figure 1:**
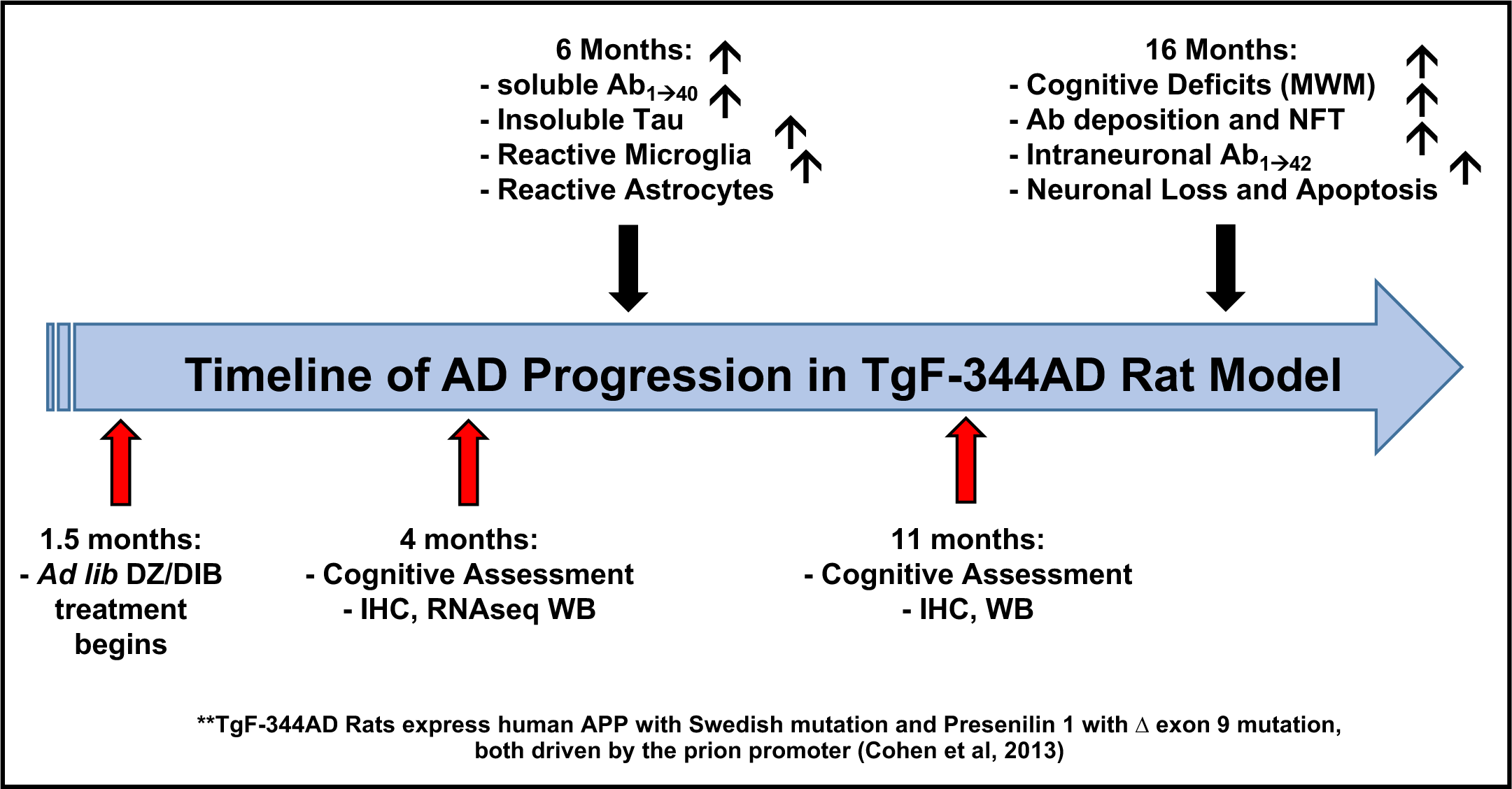
Timeline of AD progression in TgF-344 AD rats. Black arrows show development of AD hallmarks, including amyloid beta plaques, reactive microglia and astrocytes, tau tangles, cognitive deficits, neuronal loss and apoptosis. Red arrows show the intervention strategy, starting with ad lib treatment with DZ/DIB starting at 1.5 months and cognitive assessments at 4 and 11 months.

**Table 1:** The DZ/DIB Cohort is composed of 105 rats, divided among two age groups (4 and 11 months) and four treatment groups (wild type not treated, transgenic not treated, wild-type DZ/DIB treated, and transgenic DZ/DIB treatment) and collapsed across sexes (male and female).

| Treatment | Untreated |  | DZ (10mg/kg/day) + DIB (200mg/kg/day) |  |
| --- | --- | --- | --- | --- |
| Genotype | Wild Type | Transgenic-AD | Wild-Type | Transgenic-AD |
| Age | Four Months |  |  |  |
| # of Rats | 17 | 18 | 10 | 17 |
| Age | 11 Months |  |  |  |
| # of Rats | 14 | 9 | 9 | 11 |
| Total # of Rats (105 Rats) | 31 | 27 | 19 | 28 |

### 2.2 DZ/DIB treatment

Orally administered DZ/DIB treatment started when the rats were 52 days old. Tg-AD and WT rats were fed a combination of DZ (10 mg/kg bw/day, Sigma Aldrich cat# D9035) and DIB (200 mg/kg bw/day, Sigma Aldrich Cat # D33454) in rodent chow (Research Diets Inc. NJ). Non-treated Tg-AD and WT rats were fed normal chow. Treatment ended at 4- or 11-months of age when brains were collected for further analyses.

### 2.3 Radial arm maze (RAM, 8-arm)

Procedures are as previously published (Braren et al, Avila et al 2021). Briefly, rats were food-deprived to 85% of their free-feeding weight. Prior to training, rats were habituated to the maze and the food reward (*Ensure* Food Supplement) with six exposures to the maze over two days. During RAM training all 8 arms were baited with Ensure in a submerged food cup at the end of each arm. Rats received 4 training trials, two trials per day. Each trial was initiated by confining the rat to the center of the arena with an opaque covering. When the covering was removed, the rat was free to collect baits. The trial was terminated when all baits were retrieved, or 25 min had elapsed. To ensure the rats were using a spatial strategy, the maze was rotated 90 degrees each day and cleaned with 70% ethanol after each trial to prevent internal maze cues from being used.

The sequence of arm entries for each rat was recorded. Re-entry into a previously entered arm was scored as an error. We measure the number of errors for light working memory load, defined as the number of errors made collecting baits 1-4, and heavy working memory load, defined as the number of errors in collecting baits 5-8.

### 2.4 Immunohistochemistry

Hippocampal immunohistochemistry (IHC) was performed in 4- and 11-month old rats, as previously described^25,26^. Primary and secondary antibodies used are listed in supplemental Table 1. VectaShield® mounting media with DAPI (cat# H-1200-10) was used and slides were stored at 4°C in the dark until imaged.

Ramified, reactive, and amoeboid microglia phoenotypes were analyzed for circularity based on the ImageJ form factor (FF = 4π x area/perimeter^2): ramified (FF < 0.50), reactive (FF 0.50 - 0.699), and amoeboid (FF > 0.70), where microglia with an area between 50 – 1000µm^2 were included. Microglia signal intensity (O.D) was quantified as previously^25,26^.

To image the sections, we used the Zeiss AxioImager M2 wide-field fluorescence microscope combined with a Zeiss AxioCam MRm Rev. 3 camera connected to a motorized stage. We used the AxioVision 4 software, module MosaicX to capture the images. Exposure times for each channel were kept consistent across sections. Images were captured at 5x, 10x, and 20x magnifications. Captured images were saved as ZVI files, loaded onto Image J (NIH, Bethesda, MD) and converted to .tif files for use in optical density and co-localization analyses. Each channel was analyzed to an antibody specific threshold^27^.

Quantifications included the whole left hippocampus as well as hippocampal subregions CA1, CA3, dentate gyrus, and subiculum. Positive signals were measured, masks created, and merged when co-localization analyses were required. Pixel areas meeting threshold intensity criteria were measured in the delineated hippocampal regions at 10x. Pixel intensities were calculated from the images at 16-bit intensity bins. TIF files were batch processed with well-developed previously described^25,26^ scripts for each channel/marker to detect positive signals, and analyzed with GraphPad Prism 9.5 (GraphPad Software, San Diego, CA).

### 2.5 Western blot

We generated western blots of right hippocampal tissue and quantified as previously described^26^. Primary antibodies used are listed in Supplemental Table 3.

### 2.6 RNAseq analysis

We analyzed right hippocampi for gene expression by RNA sequencing (RNAseq) at the UCLA Technology Center for Genomics and Bioinformatics (Los Angeles, CA). Data were normalized as reads per million (RPM) using the TMM method. Differentially expressed genes from DZ/DIB-treated transgenic rats were determined using the edgeR program^28^. RPMs were analyzed for fold-changed, p-values, and FDR for each gene (Supplemental Table 2). A total of 35 female samples were analyzed as shown in Supplemental Table 1. We chose females because two out of three AD patients are females^29^, thus we concentrated on the female gene expression.

### 2.7 Statistical analysis

For statistical analyses, sexes were combined into four groups: WT and transgenic not treated (WTNT and TGNT) and WT and transgenic DZ/DIB treated (WTTR and TGTR) (Table 1, number of rats/group).

For behavioral and most assessments, two-way ANOVAs were used to analyze differences between genotype and treatment. Post-hoc analyses used controlled Sidak’s-corrected *t* test comparisons. One-tail independent *t*-tests were used for analysis of Aβ plaque and tau-PHF burden. For image quantification, normalization of pixel intensity values across images was done utilizing the rolling ball algorithm^30^. The macroscripts necessary for image processing and quantification were added to GitHub (https://github.com/GiovanniOliveros33/Ibudilast-Manuscript). The alpha level was set at *P* < 0.05 with a 95% confidence interval for each effect. GraphPad Prism version 9.5.1 (La Jolla, California) was used.

For western blot analysis, a mixed model three-way ANOVA was first used along with Tukey’s post-hoc analysis to measure differences in band intensity in all measured samples across two age points (4 months and 11 months), two genotypes (WT and Tg-AD), and drug treatments (DZ/DIB treatment vs. no treatment). An ordinary two-way ANOVA was also performed with Tukey’s post-hoc analysis to determine the effects of DZ/DIB treatments within each respective age group, still considering the effects of treatment and genotype on protein levels.

To measure differences in gene expression between TGNT and TGTR experimental groups multiple unpaired *t*-tests with Welch’s correction were used for each comparison. Genes considered exhibited a fold-change of at least 1.5, and an FDR of 1%, determined by the two-stage step-up method^31^. Z-scores were generated for RNAseq heat maps and were calculated with the following formula [(*Sample RPM value*) – (*Mean of all RPM values for respective gene*) / (*Standard Deviation of all RPM values for respective gene*)].

## 3. RESULTS

### 3.1 DZ/DIB treatment mitigates spatial working memory deficits exhibited by 11 months Tg-AD rats compared to WT littermates

Two-way ANOVA on light working memory load shows a significant effect of genotype and drug treatment at 11 months, Fig 2B [F_(1,39)_ = 6.09, p=0.02 for genotype; F_(1,39)_ = 12.25, p = 0.001 for drug treatment]. Sidak’s post-hoc tests show a significant difference between WTNT (n=14) and TGNT (n=9) [t=3.663; p=0.004] and between TGNT and TGTR (n=11) [t= 2.769; p = 0.05]. A similar analysis for heavy working memory load performance at 11 months showed a significant effect for genotype [F_(1,39)_ = 4.43, p = 0.04], but not for treatment (F_(1,39)_ = 0.85, p = 0.36), with no post-hoc differences (data not shown). Since 11 month old rats had also received RAM training at 4 months of age, the age effect observed in WT rats could reflect improved recall of prior training, an effect not observed in the transgenic conditions.

**Figure 2:**
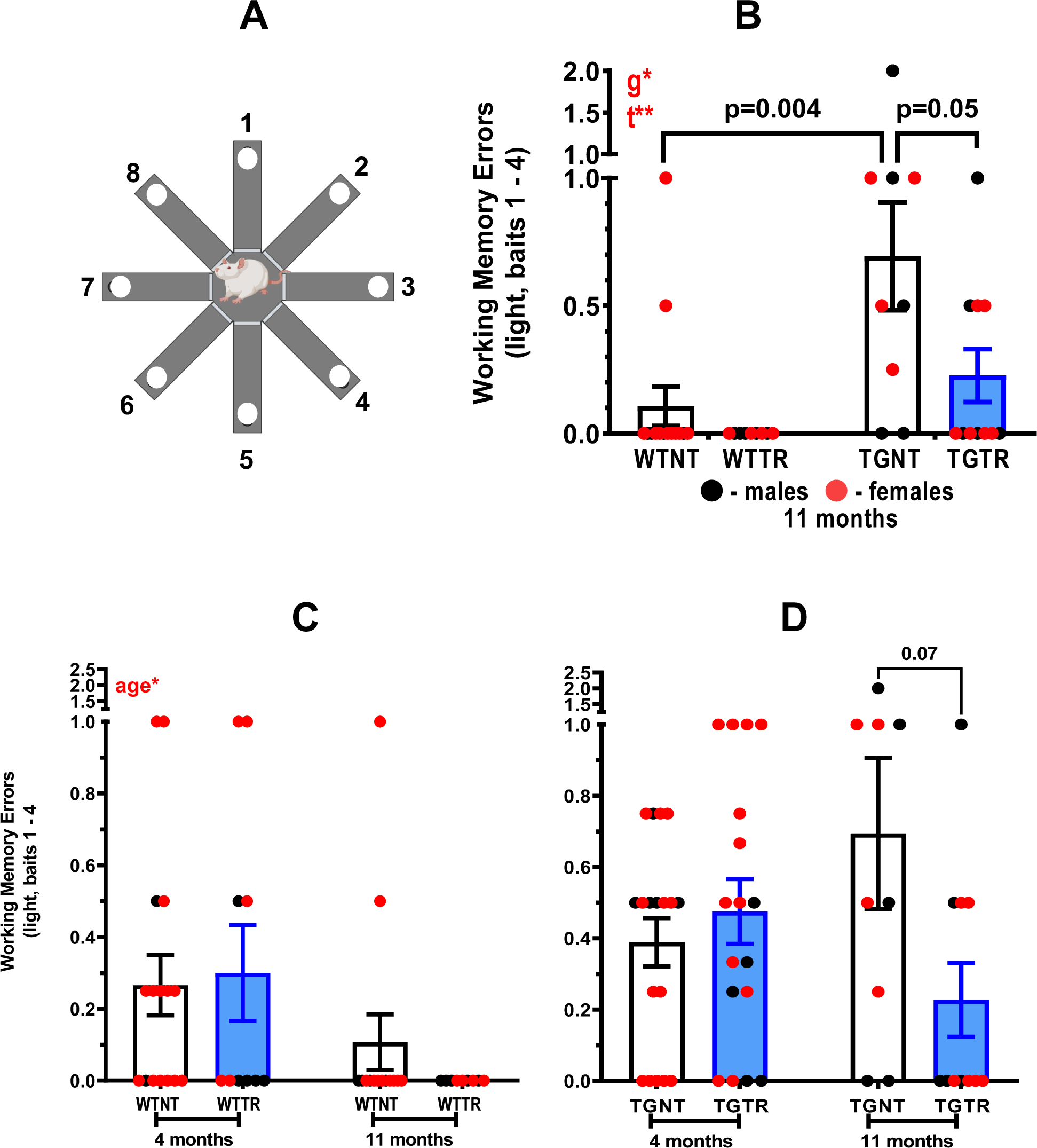
Effects of DZ/DIB treatment on cognition. (A) The 8-arm radial arm maze. (B) 11-month old Tg-AD untreated rats show increased working memory errors under a light working memory load (tasked with reaching the first four arms) when compared to Wild-type untreated rats. DZ/DIB treatment shows significant reduction in working memory errors in Tg-AD rats. (C) Age comparison among wild-type rats show that 11-month wild-type rats show improved working memory compared to 4-month wild-type rats. (D) Age-comparison among Tg-AD rats do not show any significant working memory differences between 4 and 11-month rats. *a = age effect, *g = genotype effect, *t = drug treatment effect. * = p < 0.05, ** = p < 0.01. WTNT = wild-type not treated, TGNT = transgenic-AD not treated, WTTR = wild-type DZ/DIB treated, TGTR = transgenic-AD DZ/DIB treated.

### 3.2 DZ/DB does not enhance memory between 4 month and 11 month old rats

At 4-months of age WT rats performed significantly worse on the light working memory load assessment than when retested at 11 months of age, independently of treatment [Figure 2C, F_(1-45)_ = 6.27, p = 0.02 for age, no post-hoc differences; F_(1-45)_ = 0.16, p = 0.69 for drug treatment]. In contrast, there were no significant differences between the light working memory performance of transgenic 4-month compared to 11-month rats [Figure 2D, F_(1-51)_ = 0.07, p = 0.80 for age; F_(1-51)_ = 2.89, p = 0.10 for drug treatment]. This could indicate that the 11-month WT rats remembered the task performed when they were first exposed to RAM at 4-months, but the transgenic rats did not. A similar two-way analysis of heavy working memory load comparing 4-month and 11-month rats shows an effect of age [F_(1,54)_ = 16.45, p < 0.001 for WT; F_(1,43)_ = 4.96, p = 0.03 for transgenic], with 4-month old rats performing better than 11-month old rats. No significant drug treatment differences were observed for WT [F_(1,54)_ = 0.008, p = 0.93], or transgenic rats [F_(1,43)_ = 0.02, p = 0.88] (data not shown).

### 3.3 DZ/DIB treatment significantly reduces A**β** plaque burden in 11-month Tg-AD rats compared to untreated Tg-AD littermates

DZ/DIB treatment significantly reduced Aβ plaque burden (red) in the hilar subregion of the DG in the left hippocampus of 11-month TGTR (Figure 3B, D) compared to TGNT rats (Figure 3A, C). The difference is statistically significant (Figure 3E, t = 2.140, df = 17, p = 0.05). DZ/DIB treatment did not affect Aβ plaque buildup throughout the whole left hippocampus and in subregions CA1, CA3, and the SB of 11-month Tg-AD rats compared to Tg-AD untreated littermates (Supplemental Table 1). In addition, 4-month-old TG-AD rats or WT rats of both ages did not develop Aβ plaques independently of drug treatment (not shown).

**Figure 3:**
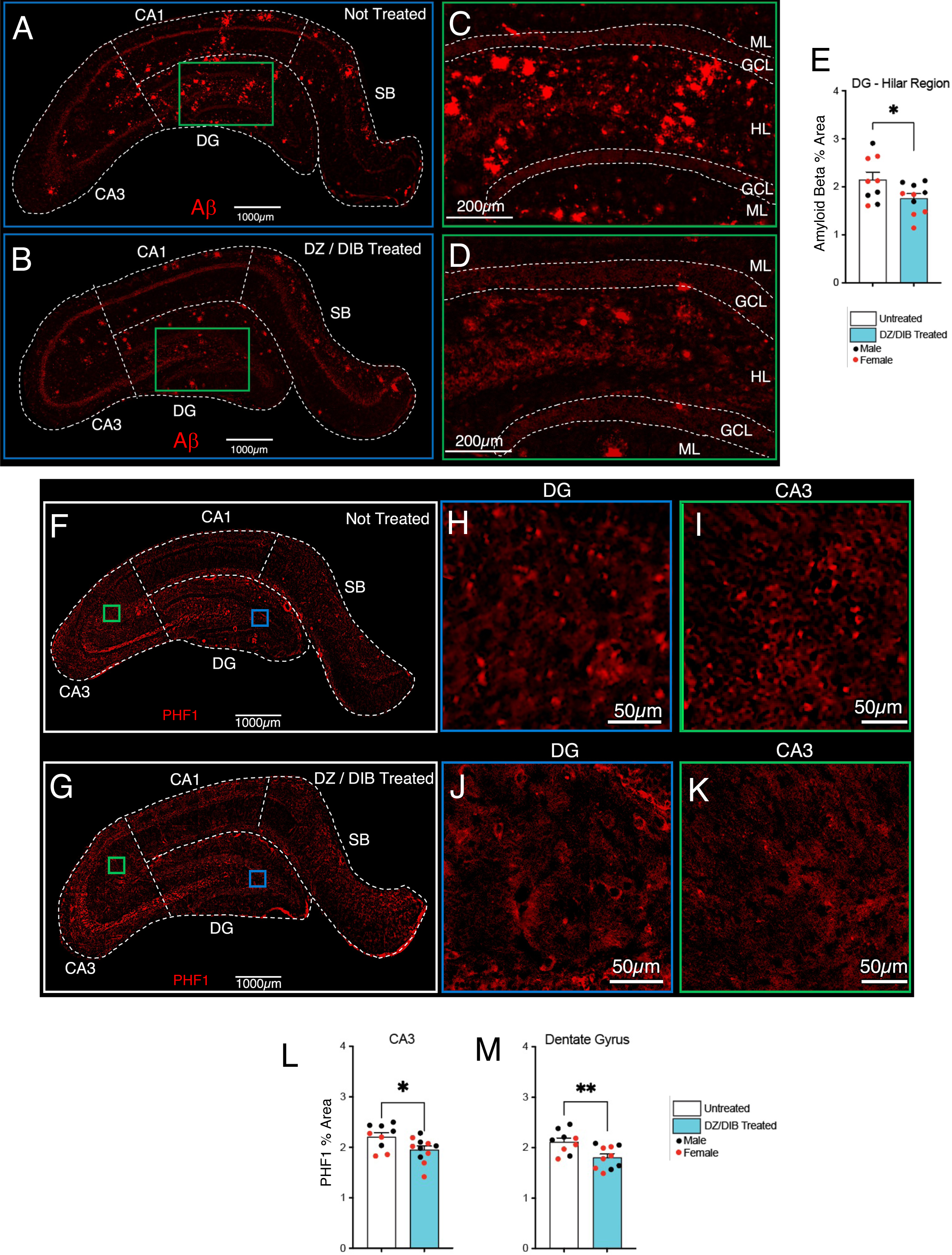
Effects of DZ/DIB Treatment on Alzheimer’s disease hallmarks. 11 month-old Tg-AD rats stained by IHC for Amyloid beta plaques across hippocampal regions for Tg-Not Treated (A) and Tg-DZ/DIB Treated (B) rats. Scale bar = 1000µm for large panels (A and B). Dentate Gyrus (DG) magnifications for Tg-Not Treated (C) and Tg-DZ/DIB Treated Rats (D). Scale bar = 200µm for magnification of dentate gyrus panels (C and D). Within the dentate gyrus (C and D), analysis of the hilar region of DG showed DZ/DIB treatment reduced amyloid beta levels compared to untreated rats (E). 11 month-old Tg-AD rats stained by IHC for Tau paired helical filaments (PHF1), an early precursor of neurofibrillary tangles across hippocampal regions for Tg-Not Treated (F, H, and I) and Tg-DZ/DIB Treated rats (G, J and K). Scale bar = 1000µm for large panels (F and G). Scale Bar = 50µm for magnification of dentate gyrus panels (H and J), and CA3 panels (I and K). 11 month DZ/DIB treated rats exhibited significant reductions in PHF1 levels in CA3 (L) and the dentate gyrus (M). Unpaired two-tailed t (-tests with Welch’s corrections were used for analyses performed in panels E, L and M. *p < 0.05, **p < 0.05. n = 10 TGNT, n = 10 TGTR. Tg-AD = Transgenic – Alzheimer’s disease. CA = cornu ammonis, DG = dentate gyrus, SB = subiculum, PHF = paired helical filaments, ML = molecular layer, GCL = granular cell layer, HL = hilar region.

### 3.4 DZ/DIB treatment significantly reduces tau PHF1 levels in the hippocampal DG and CA3 subregions of 11-month Tg-AD rats compared to untreated Tg-AD littermates

DZ/DIB treatment reduced PHF1 levels in the DG and CA3 hippocampal subregions of TGTR (Figure 3G, J, and K) compared to TGNT (Figure 3F, H, and I) rats. The decrease in PHF1 levels was significant in the CA3 (Figure 3L, t = 2.30, df = 17.06, p = 0.034) and DG (Figure 3M, t = 2.96, df = 16.7, p = 0.009) subregions. However, no significant reductions in PHF1 signal were detected in CA1 and SB (Supplemental Table 2) subregions upon DZ/DIB treatment. We did not analyze tau PHF1 levels in 4-month-old Tg-AD rats since prior reports showed no accumulation of PHF1 until later ages^32^.

### 3.5 DZ/DIB treatment did not significantly impact neuronal loss and microgliosis exhibited by Tg-AD rats compared to their WT littermates

At 4 months of age there were no significant hippocampal neuronal losses or increases in microglia numbers in TGNT compared to WTNT rats.. In contrast, by 11 months of age, there were significant hippocampal neuronal losses particularly in the DG granular cell layer (GCL) in TGNT compared to WTNT rats (Supplemental Table 3A and 3B). Moreover, hippocampal total microglia, as well as ramified, reactive, and amoeboid microglia considered separately were increased in Tg-AD compared to WT rats. (Supplemental Table 4A through 4H). The latter types of microglia represent the three distinct microglia morphologies, based on their cell body shape and number of processes and branching^33^. Overall, DZ/DIB treatment did not mitigate neuronal loss or increased levels of microglia detected in the 11-month Tg-AD rats compared to the WT littermates.

### 3.6 APP levels are increased in 4-month transgenic rats following DZ/DB treatment

We assessed the levels of full-length APP by western blot using the mouse monoclonal antibody 22C11 which detects both human and rat APP (Figure 5A). A three-way ANOVA analysis of APP levels across genotype show Tg-AD rats express higher levels of APP compared to age-matched WT rats [*F*_(1, 39)_ = 98.05; p < 0.0001] (not shown). Across age however, non-treated 11-month Tg-AD rats express higher APP levels than 4-month Tg-AD rats [*F*_(1, 39)_ = 12.49; p = 0.0011]. In addition, DZ/DIB treatment increases APP levels compared to untreated rats, irrespective of genotype (F_(1,39)_ = 8.63, p = 0.006). Significant post-hoc differences were observed between 4 month WT untreated and 4 month TG untreated rats (p = 0.0236), between 4 month TG untreated and 11 month TG untreated rats (p = 0.0149), between 11 month WT untreated and 11 month TG untreated rats (p < 0.0001), and between 11 month WT DZ/DIB treated and 11 month TG DZ/DIB treated rats (p = 0.0024). No other significant post-hoc differences were observed in the 3-way ANOVA analysis.

**Figure 4:**
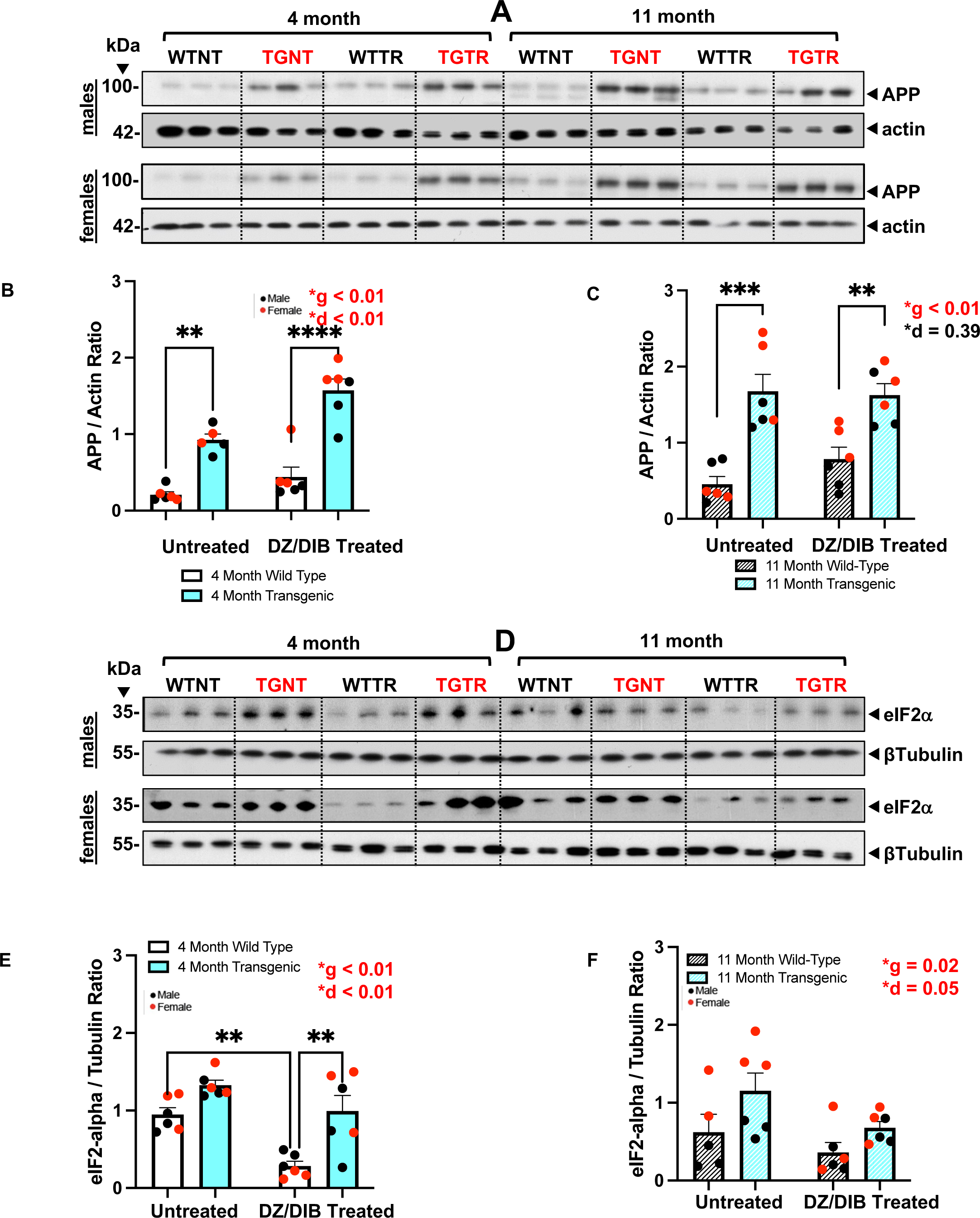
DZ-DIB Treatment effects on APP and eIF2-alpha levels. Male and Female rat hippocampal tissue homogenates were used to measure DZ-DIB treatment effects on full-length APP (A) and eIF2-alpha (D) levels by western blot analysis. Data represents the percentage of the pixel ratio for full-length APP and eIF2-alpha over their respective loading controls (actin for APP and beta-tubulin for eIF2-alpha). Analysis was performed for the four groups WTNT, WTTR, TGNT, TGTR at 4 months (B for APP, E for eIF2-alpha) and 11 months (C for APP and F for eIF2-alpha). All graphed data represent the ratio either APP or eIF2-alpha over their respective loading controls (Actin for APP and Tubulin for eIF2-alpha). Values are means + SEM. Significance (asterisks shown on graphs) represent post-hoc effects following an ordinary two-way ANOVA with Sidak’s post-hoc tests. *p < 0.05, **p < 0.01, ***p < 0.001, ****p < 0.0001. n = 6 WTNT, n = 6 WTTR, n = 6 TGNT, n = 6 TGTR. WTNT = Wild-type not treated, TGNT = transgenic not treated, WTTR = wild-type DZ/DIB treated, TGTR = transgenic DZ/DIB treated. g = genotype effect, d = DZ/DIB drug treatment effect.

**Figure 5:**
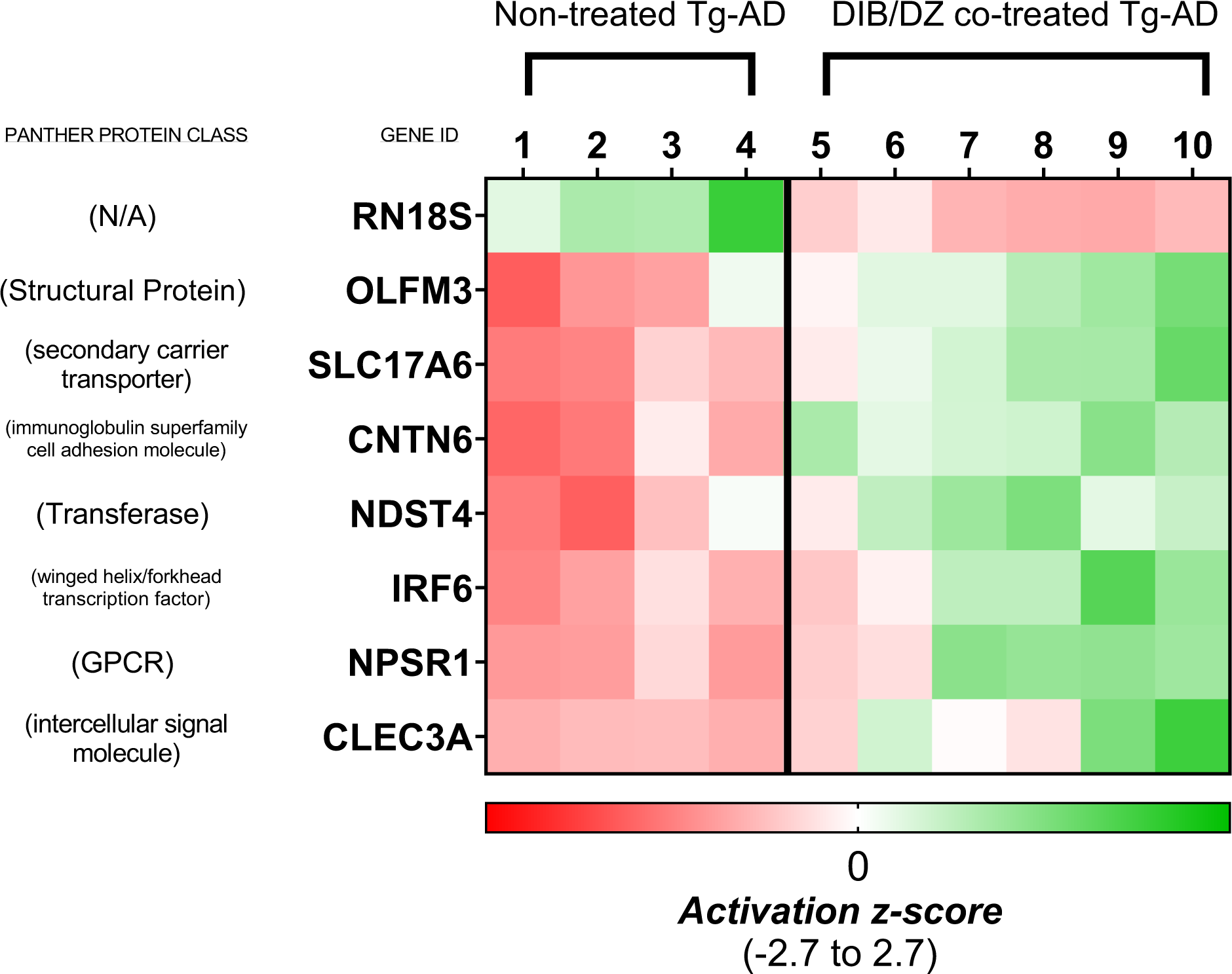
RNAseq heatmap shows transcriptional changes in Tg-AD rats following DZ/DIB treatment. Treatment with DZ/DIB shows downregulation in RN18S, while showing increased regulation of OLFM3, SLC17A6, CNTN6, NDST4, IRF6, NPSR1, and CLEC3A. Changes in expression are represented as changes in activation z-score (ranging from -2.7 to 2.7). n = 4 Non-treated Tg-AD rats, n = 6 DZ/DIB co-treated Tg-AD rats.

Further two-way ANOVA analysis at 4-months revealed significant genotype differences (*F*_(1, 19)_ = 71.91; p < 0.001) in which Tg-AD AD rats had higher levels of APP regardless of drug treatment (Figure 4B). Drug treatment effects were also observed, where DZ/DIB treatment showed high levels of APP compared to untreated rats at 4 months (*F*_(1,19)_ = 16.30, p < 0.01). Post-hoc analysis showed differences between 4-month WT and 4-month Tg-AD untreated rats (p < 0.01), and between 4-month WT DZ/DIB treated rats and 4-month Tg-AD DZ/DIB treated rats (p < 0.01). At 11-months, significant genotype differences (*F*_(1, 20)_ = 40.38; p < 0.001) were observed, in which Tg-AD rats had higher levels of APP regardless of drug treatment. (Figure 4C). Post-hoc analysis showed differences between 4-month WT and 4-month Tg-AD untreated rats (p < 0.01), and between 4-month WT DZ/DIB treated rats and 4-month Tg-AD DZ/DIB treated rats (p < 0.01).

### 3.7 DZ/DIB treatment attenuates the increases in eIF2**α** levels detected in Tg-AD rats compared to WT littermates

The translational control factor eIF2α along with its phosphorylating kinases such as PERK, act as a critical molecular switch that inhibits global protein translation during the unfolding protein response (UPR)^16^. This molecular switch is overactive in AD and thus, attenuating the activity of this molecular switch would be beneficial to AD.^16,34^ We assessed the levels of eIF2α by western blot (Figure 4D). A three-way ANOVA analysis revealed an overall attenuating effect of DZ/DIB treatment on eIF2α levels irrespective of age and genotype [F_(1, 39)_ = 17.06; p = 0.0002] (not shown). Interestingly, eIF2α levels were increased in Tg-AD rats compared to WT littermates, independently of age and treatment [F_(1, 39)_ = 21.45; p < 0.0001]. No age specific effects were observed [*F*_(1, 39)_ = 3.09; p = 0.086].

Further analysis for age-specific effects were examined with an ordinary two-way ANOVA that showed at 4 months, Tg-AD rats had higher levels of eIF2α compared to WT littermates (F_(1, 20)_ = 20.94; p = 0.0002), and DZ/DIB drug treatment attenuated the levels of eIF2α (F_(1, 20)_ = 17.61; p = 0.0004) regardless of genotype (Figure 4E). Significant post-hoc effects were observed between untreated wild type and DZ/DIB treated wild-type rats (p = 0.004), and between DZ/DIB wild-type and DZ/DIB transgenic rats (p = 0.002). At 11 months (Figure 5F), transgenic rats had higher eIF2a levels compared to wild-type rats, irrespective of drug treatment (F_(1,19)_ = 5.949, p = 0.0247). In addition we observed an overall drug effect, where DZ/DIB attenuated eIF2-alpha levels irrespective of genotype (F_(1,19)_ = 4.396, p = 0.0496). No significant post-hoc differences were observed.

### 3.8 Transcriptome changes in DZ/DIB treated compared to untreated 4-month Tg-AD female rats

Approximately two thirds of all Alzheimer’s patients are women. Moreover, there is a critical need to identify early biomarkers, for their ability to indicate early stages of AD when treatment can be most effective. Thus, compared the transcriptomes of 4-month untreated and treated Tg-AD female rats. Differences in the expression of the APP and PSEN1 genes between untreated WT and Tg-AD was expected, because the Tg-AD rats express APPswe and PS1ΔE9 mutations driven by the prion promoter, at 2.6- and 6.2-fold higher levels than the respective endogenous rat protein. The RNAseq showed that APP and PSEN1 are upregulated at both 4 and 11 months (Table 4.1 A and B).

For the treatment, the RNAseq data also showed a ∼2.0 fold decrease in EGR2 and a ∼3.9 fold increase in HIST1H2AA expression in female TGNT rats compared to WTNT at 4-months of age (Table 4.1A). There was a ∼2.0 fold increase in GFAP and a ∼182.1 fold increase in RN7SL1 expression in female TGNT rats compared to WTNT at 11-months of age (Table 4.1B). TGTR rats saw an increased fold change in expression for eight genes: OLFM3 (∼1.6), CNTN6 (∼1.8), NDST4 (∼1.9), IRF6 (∼2.8), SLC17A6 (∼1.8), KL (∼31.3), NPSR1 (∼4.2) and CLEC3A (∼15.4) compared TGNT at 4 months (Table 4.1C). RN18S was decreased ∼1.6 fold in TGTR rats compared TGNT at 4 months (Table 4.1C).

## Sex-dependent changes in gene expression in treated Tg-AD rats

The RNAseq data showed a ∼2 fold decrease in EGR2 expression in female TGNT rats at 4 months of age. EGR2, is an inducible transcription factor and being that this is the only transcriptionally downregulated protein in TGNT rats we investigated the expression by WB. EGR proteins have long been studied in the CNS for their role subtly controlled changes in gene expression^35^. Studies on EGR2 suggest that EGR2 plays a role in regulating the activity of NF-κB and deletion of EGR2 is lethal in mice ^36,37^.

## Discussion

Our study investigated the repurposing potential for DZ and DB as a combination treatment for AD. DZ has been used for decades for its cardioprotective effects, showing it is a well-tolerated and efficacious drug, therefore a great candidate for repurposing ^10^. DZ’s mechanism against apoptosis may be helpful in preventing neuronal loss in later stages of AD, but it was not robustly seen at 11-months of age. DIB is a promiscuous drug with affinity for many targets and likely some still unknown. Some of the suggested mechanisms for DIB are inhibition of the unfolded protein response, increase in activity of protective transcription factors, induction of cell cycle arrest, and reducing expression of androgen receptors ^16,23,38^. DIB was shown to be neuroprotective to hippocampal histopathology^16^, but to our knowledge, not in an AD animal model.

Our data show that DZ/DIB cotreatment reduced Aβ plaque burden, reduced hyperphosphorylated tau, and decreased eIF2α levels in the hippocampus. However, DZ/DIB cotreatment did not reduce gliosis, prevent GCL neuronal loss, or alter EGR2 levels. The cognitive assessment at 11-months of age shows that DZ/DB treatment mitigated spatial deficits in light working memory load performance in TGTR vs TGNT and there was an overall treatment benefit across all groups indicating an effect beyond genotype. Additionally, there was an age-dependent decline in Heavy Working Memory Load irrespective of genotype likely because this portion of RAM becomes increasingly challenging where we can see age related decline. Interestingly analysis of 4 month vs 11 month performance on light working memory load, shows that WT rats performed better at 11-months compared to 4-months of age. This is likely due to recall of prior exposure to the task, an effect not observed in the transgenic conditions.

We show enhanced levels of Aβ plaque and neurofibrillary tau tangles in the DG hilar subregion in transgenic animals and subsequent plaque reduction with DZ/DIB treatment, indicating that the DG is more susceptible to insults and more prone to develop plaques.^39^ This same susceptibility of the DG region to development of AD pathology earlier than other hippocampal subregions may explain why at 11 months of age, the neuronal loss exhibited in Tg-AD rats is limited to the DG region, particularly in the granular cell layer, consistent with prior reports.^25^

DZ/DIB treatment induced a decrease in the levels of eIF2-α protein in the overall hippocampus. eIF2-α is a subunit within the eIF2 initiation complex responsible for the regulation of global and specific mRNA translation^16^. Phosphorylation of eIF2-α by stress-activated kinases (such as pERK, HIR and KPR) prevents nucleotide exchange and sequesters eIF2B, limiting the availability of eIF2-α/GDP complex, leading to decreases in translation. This suggests that DZ/DB reduces the likelihood for premature translational depression, consistent with prior DIB treatment investigations^16^.

DZ/DB treatment reduced PHF1 levels extending beyond the hilar region encompassing the entire DG regions as well as CA3. Formation of neurofibrillary tau tangles are suggestive of neuronal cell stress due to disruptions to microtubule structure caused by hyperphosphorylation of tau^40^. Thus it is plausible that lowering eIF2-α levels with DZ/DIB treatment may make neuronal cells less prone to the formation of PHFs, the precursors to tau tangles short-term due to continued neuronal cell protein synthesis.

RNAseq analysis between female TGNT and WTNT show significant changes at 4-months of age in only two genes: Histone H2A type 1-A (*HIST1H2AA*) and Early Growth Response 2 (*EGR2)*. *HISIT1H2AA* is a subtype of the H2A histone which is one of five histones that package DNA to make up the nucleosome important for gene regulation ^41^. Histones regulate gene expression, and they are post-transcriptionally modified with mechanisms such as acetylation, methylation, phosphorylation and ubiquitination^42^. A study using human frontal cortex brain tissue found that H2A was less ubiquitinated in AD brain tissue compared to age matched controls ^43^, so the increase in *HISIT1H2AA* could result from a reduction in ubiquitin/proteasome degradation. Further studies would be required to conclude if this upregulation is a result of less ubiquitin/proteasome degradation.

As expected, the RNAseq analysis shows that the TgAD rat overexpresses Amyloid Precursor Protein (*APP)* and Presenilin 1 (*PSEN1)* at both 4- and 11-months of age, as well as *GFAP* being upregulated only at 11-months of age indicating astrogliosis with increased neuroinflammation. The highest fold difference in the RNAseq that was observed at 11-months of age between the TGNT and WTNT rats was the RNA component of the signal recognition particle 7SL1 (*RN7SL1*). *RN7SL1* is an important component in targeting proteins for transport throughout the cell, elevated levels of *RN7SL1* leads to increased inflammation^44,45^. Thus, the 182-fold increase in TGNT rats may be due to impaired transport or neuroinflammation.

The RNAseq analysis in TGTR vs TGNT rats was performed at 4 months of age to assess how the DZ/DIB treatment may alter gene expression in early AD pathogenesis. Genes that were upregulated are connected to cognition, neurogenesis, differentiation, synaptic plasticity, apoptosis, and amyloid toxicity. The genes that were up regulated in TGTR vs TGNT rats were: (1) <u>Olfactomedin 3</u> (*OLFM3*), which is important for differentiation in the brain and retina ^46^. (2) <u>Contactin 6</u> (*CNTN6*) that plays a role in axonal formation for the mossy fibers within the hippocampus during development and it also is thought to be involved in apoptosis for neuron survival ^47^. (3) <u>N-deacetylase and N-sulfotransferase 4</u> (*NDST4*) identified a markers for recently differentiated neural cells ^48^. (4) <u>Interferon regulatory factor 6</u> (*IRF6*) which is relevant to epithelial tissue, where it mediates differentiation and helps regulate apoptosis ^49,50^. (5) <u>Solute</u> <u>Carrier Family 17 Member 6</u> (*SLC17A6*) that enables the activity of glutamate transporters and is important for synaptic plasticity and expression correlated to cognitive function ^51,52^. (6) <u>Neuropeptide S receptor 1</u> (*NPSR1*) is important in hippocampal function and promotes synaptic plasticity ^53^. (7) <u>Rn18s 18S ribosomal RNA</u> (*RN18S*) and (8) <u>C-type lectin domain family 3</u> <u>member A</u> (*CL3C3A*) induce changes in transcription that are less clear, because respectively one has ribosomal function while the other is known to bind to heparin ^54,55^. RN18S was the only gene that was not identified by PANTHER’s Gene Ontology Tool, so it is difficult to hypothesize the relevance. For the two excluded proteins due to variability, *TTR* which is a multifunctional protein that has many roles in the CNS including cognition and neurogenesis via interaction and transport of thyroxine and retinol (Vitamin A) [17, 18] and has been shown to be neuroprotective against AD via binding to Aβ40 to prevent Aβ toxicity [19], and *KL* has been shown to mitigate amyloid burden derived from APOE4 pathways ^56^.

The following upregulated genes after DIB/DZ treatment are known to decline with AD severity and/or ageing: SLC17A6, TTR, KL and NDST4 ^52,57-59^. It is possible that the pathology that contributes to a decline in the transcriptome of the TGNT may have been mitigated by the DZ/DIB treatment at 4-months of age. Very important to note, the single drug treatments of DIB and DZ had no effect on the transcriptome of Tg-AD rats whereas the combination of DZ/DIB did. The long-term effects of treatment on these proteins during moderate pathology is unknown and should be explored at later disease phenotypes.

Our treatment covered a wide range of targets including common markers for AD as well as specific targets for DZ and DIB. DZ/DIB treatment showed reductions in Aβ plaque and tau PHF1 loads. This is consistent with prior investigations using the individual drugs.^25^ DZ/DIB treatment could be maintaining protein translation and showing some neuroprotective potential due to reductions in eIF2α expression upon treatment. Treatment of these two drugs widens the reach of polypharmacology, with multiple mechanisms of action such as DZ balancing calcium overload, DIB’s relief of cellular stresses through eIF2 and activation of nuclear factor erythroid 2–related factor 2 (Nrf2), and inhibition of c-Jun N-terminal kinases (JNK) to prevent oxidative damage. DZ increases intracellular potassium levels and conversely reduces intracellular calcium levels, while DIB has a host of effects on signaling pathways involved in cancer, mitigating inflammation and cellular stress, and neuroprotection.^18^

Although current FDA approved AD therapeutics are managing symptoms to varying degrees of success, they currently do not address biological changes that initiate the cascade of changes observed in disease pathology and progression. A likely culprit is the multifactorial nature of AD, which multi-drug therapies have the potential to address through targeting of multiple signal pathways and mechanisms that contribute to AD progression. Shifting efforts to computational models and polypharmacology is on the rise and can prove the be an invaluable asset, to which our findings have come to suggest.

