## Supplemental Figures for "Diazoxide/dibenozylmethane treatment mitigates spatial memory deficits and pathology and upregulates protective genes in an Alzheimer’s transgenic rat model"

**Supplemental Table 1:** Amyloid Beta (Aβ 4G8) percent positive signal across hippocampal regions at 11 months of age

**Tg-AD untreated	Tg-AD DZ / DIB**

**Region	mean ± SEM	treated	*p*-value	*t*-statistics**

**mean ± SEM**

HC	1.785 + 0.10	1.777 + 0.10	0.94	t = 0.080, df = 17.00

SB	1.979 + 0.18	2.053 + 0.18	0.70	t = 0.397, df = 17.00

CA1	2.019 + 0.26	2.441 + 0.42	0.12	t = 1.611, df =17.00

CA3	1.404 + 0.04	1.447 + 0.14	0.77	t = 0.304 , df = 17.00

DG	1.922 + 0.15	1.774 + 0.12	0.26	t = 1.159 , df = 17.00

**Supplemental Table 1: Amyloid Beta (Aβ 4G8) levels across the hippocampus in 11-month Tg-AD rats**. Values represent the percent positive signal detected within a specific hippocampal area (out of 100%) as explained in materials and methods for 11-month rats. Data was analyzed using unpaired one-tailed t-tests with Welch’s corrections. Abbreviations: HC – hippocampus: CA – cornu ammonis: DG – dentate gyrus: SB – subiculum: Tg-AD – transgenic model of Alzheimer’s disease: DZ / DIB: Diazoxide / Dibenozylmethane. SEM: standard error of the mean. Tg-AD untreated (11 months): n = 9, Tg-AD DZ/DIB treated (11 months): n = 10.

**Supplemental Table 2:** Tau neurofibrillary tangle (PHF1) percent positive signal across hippocampal regions at 11 months of age

**Tg-AD untreated	Tg-AD DZ / DIB**

**Region	mean ± SEM	treated	*p*-value	*t*-statistics**

**mean ± SEM**

HC	0.820 + 0.06	0.756 + 0.05	0.19	t = 1.358, df = 16.84

SB	1.590 + 0.15	1.745 + 0.23	0.52	t = 0.666, df = 14.86

CA1	2.211 + 0.06	2.273 + 0.18	0.74	t = 0.338, df =17.82

CA3	2.209 + 0.26	1.954 + 0.11	0.03	t = 2.299 , df = 17.06

DG	2.115 + 0.31	1.807 + 0.10	< 0.01	t = 2.957 , df = 16.67

**Supplemental Table 2: Tau (PHF1) neurofibrillary tangle levels across the hippocampus in 11-month Tg-AD rats**. Values represent the percent positive signal detected within a specific hippocampal area (out of 100%) as explained in materials and methods for 11-month rats. Data was analyzed using unpaired one-tailed t-tests with Welch’s corrections. Abbreviations: HC – hippocampus: CA – cornu ammonis: DG – dentate gyrus: SB – subiculum: Tg-AD – transgenic model of Alzheimer’s disease: DZ / DIB: Diazoxide / Dibenozylmethane. SEM: standard error of the mean. Tg-AD untreated (11 months): n = 9, Tg-AD DZ/DIB treated (11 months): n = 10.

**Supplemental Table 3A:** NeuN (neuronal marker) percent positive signal across hippocampal regions at 4 months of age

**WT untreated	Tg-AD untreated mean**

**Region	mean ± SEM	± SEM	*p*-value	*t*-statistics**

HC	19.94 + 0.49	19.43 + 0.55	0.25	t = 0.682, df = 13.83

SB	17.42 + 0.32	16.27 + 0.64	0.07	t = 1.592, df = 10.26

CA1	16.48 + 0.45	15.67 + 0.64	0.16	t = 1.029, df =12.63

CA3	16.57 + 0.27	16.41 + 0.28	0.34	t = 0.401 , df = 13.98

DG	17.71 + 0.47	18.25 + 0.37	0.19	t = 0.901 , df = 13.28

**Supplemental Table 3B:** NeuN (neuronal marker) percent positive signal across hippocampal regions at 11 months of age

| **Region** | **WT Untreated mean + SEM** | **Tg-AD untreated mean + SEM** | **WT DZ/DIB treated**  **mean + SEM** | **Tg-AD DZ/DIB treated**  **mean + SEM** | **p-value** | **F(DFn, DFd)** |
| --- | --- | --- | --- | --- | --- | --- |
| Hippocampus | 20.30 + 0.86 | 17.08 + 0.56 | 19.53 + 0.54 | 16.48 + 0.70 | < 0.01 (g)  0.33 (d) | F (1,33) = 20.34 (g)  F (1,33) = 0.96 (d) |
| SB | 18.51 + 0.60 | 16.58 + 0.69 | 18.10 + 0.48 | 14.98 + 0.61 | < 0.01 (g)  0.11 (d) | F (1,34) = 17.36 (g)  F (1,34) = 2.76 (d) |
| CA1 | 17.56 + 1.20 | 16.53 + 0.77 | 17.71 + 0.68 | 14.63 + 0.86 | 0.03 (g)  0.35 (d) | F (1,35) = 0.90 (g)  F (1,35) = 5.00 (d) |
| CA3 | 16.69 + 0.96 | 15.66 + 0.55 | 15.84 + 0.53 | 14.46 + 0.46 | 0.06 (g)  0.11 (d) | F (1,35) = 3.69 (g)  F (1,35) = 2.68 (d) |
| DG | 19.21 + 0.83 | 17.23 + 0.58 | 19.17 + 0.63 | 16.87 + 0.58 | < 0.01 (g)  0.77 (d) | F (1,35) = 10.14 (g)  F (1,35) = 0.09 (d) |
| GCL  (GCL / DG Area) | 4.57E-03 + 1.10E-03 | 3.77E-03 + 9.52E-04 | 4.37E-03 + 1.74E-03 | 3.81E-03 + 6.38E-04 | < 0.01 (g)  0.51 (d) | F (1,35) = 35.92 (g)  F (1,35) = 0.45 (d) |

**Supplemental Table 3: Mature neuronal cell detection in the hippocampus with NeuN**. Values represent the percent positive signal detected within a specific hippocampal area (out of 100%) as explained in materials and methods for 4-month rats (Table 1A) and 11-month rats (Table 1B). For 11-month granular cell layer measurements, data is represented as the ratio of percent positive signal in the granular cell layer divided by the total area of the dentate gyrus (total area of positive signal in GCL divided by total area of the DG). Data in table 1A was analyzed using unpaired one-tailed t-tests with Welch’s corrections. Data in table 1B was analyzed using an ordinary two way ANOVA with Sidak’s post-hoc tests. Abbreviations: HC – hippocampus: CA – cornu ammonis: DG – dentate gyrus: SB – subiculum: GCL – granular cell layer: Tg-AD – transgenic model of Alzheimer’s disease: WT – Wild-type: DZ / DIB: Diazoxide / Dibenozylmethane. SEM: standard error of the mean. WT (4 months) n = 8: TG (4 months) n = 8: WT untreated (11 months): n = 10, Tg-AD untreated (11 months): n = 10, Wild-type DZ/DIB treated (11 months): n = 10, Tg-AD DZ/DIB treated (11 months): n = 10. (g) = genotype; (d) = drug treatment.

**Supplemental Table 4A:** Total Microglia Density (represented as count per square micron) across hippocampal regions at 4 months

**WT untreated	Tg-AD untreated mean**

**Region	mean ± SEM	± SEM	*p*-value	*t*-statistics**

HC	2.79E-04 ± 5.02E-06	2.79E-04 ± 9.13E-06	0.46	t = 0.082, df = 10.88

SB	2.48E-04 ± 7.09E-06	2.51E-04 ± 9.45E-06	0.42	t = 0.218, df = 12.99

CA1	2.80E-04 ± 4.79E-06	2.79E-04 ± 1.13E-05	0.47	t = 0.069, df = 9.42

CA3	2.93E-04 ± 7.64E-06	2.93E-04 ± 1.06E-05	0.49	t = 0.018, df = 12.76

DG	2.94E-04 ± 4.35E-06	2.89E-04 ± 9.93E-06	0.32	t = 0.472, df = 9.59

**Supplemental Table 4B:** Total Microglia Density (represented as count per square micron) across hippocampal regions at 11 months

| **Region** | **WT Untreated mean + SEM** | **Tg-AD untreated mean + SEM** | **WT DZ/DIB treated**  **mean + SEM** | **Tg-AD DZ/DIB treated**  **mean + SEM** | **p-value** | **F(DFn, DFd)** |
| --- | --- | --- | --- | --- | --- | --- |
| Hippocampus | 2.53E-04 + 4.52E-06 | 3.16E-04 + 2.24E-05 | 2.58E-04 + 5.86E-06 | 3.06E-04 + 1.43E-05 | < 0.01 (g)  0.86 (d) | F (1,34) = 17.08 (g)  F (1,34) = 0.03 (d) |
| SB | 2.40E-04 + 5.85E-06 | 3.01E-04 + 2.22E-05 | 2.31E-04 + 1.02E-05 | 2.97E-04 + 1.61E-05 | < 0.01 (g)  0.68 (d) | F (1,34) = 18.57 (g)  F (1,34) = 0.17 (d) |
| CA1 | 2.59E-04 + 4.17E-06 | 3.27E-04 + 2.25E-05 | 2.65E-04 + 2.90E-06 | 3.28E-04 + 1.45E-05 | < 0.01 (g)  0.78 (d) | F (1,34) = 23.70 (g)  F (1,34) = 0.08 (d) |
| CA3 | 2.67E-04 + 6.11E-06 | 3.17E-04 + 5.00E-06 | 2.70E-04 + 1.90E-05 | 3.00E-04 + 1.20E-05 | < 0.01 (g)  0.54 (d) | F (1,34) = 11.67 (g)  F (1,34) = 0.38 (d) |
| DG | 2.57E-04 + 4.62E-06 | 3.26E-04 + 2.72E-05 | 2.60E-04 + 4.00E-06 | 3.14E-04 + 1.61E-05 | < 0.01 (g)  0.77 (d) | F (1,34) = 15.15 (g)  F (1,34) = 0.08 (d) |

**Supplemental Table 4C:** Ramified Microglia Density (represented as count per square micron) across hippocampal regions at 4 months

**WT untreated	Tg-AD untreated mean**

**Region	mean ± SEM	± SEM	*p*-value	*t*-statistics**

HC	2.62E-04 ± 8.60E-06	2.62E-04 ± 4.71E-06	0.48	t = 0.047, df = 10.84

SB	2.31E-04 ± 9.17E-06	2.22E-04 ± 4.96E-06	0.22	t = 0.805, df = 10.77

CA1	2.67E-04 ± 1.07E-05	2.69E-04 ± 5.00E-06	0.42	t = 0.197, df = 9.90

CA3	2.78E-04 ± 1.09E-05	2.83E-04 ± 7.11E-06	0.34	t = 0.414, df = 12.04

DG	2.73E-04 ± 9.10E-06	2.78E-04 ± 5.72E-06	0.35	t = 0.407, df = 11.79

**Supplemental Table 4D:** Ramified Microglia Density (represented as count per square micron) across hippocampal regions

| **Region** | **WT Untreated mean + SEM** | **Tg-AD untreated mean + SEM** | **WT DZ/DIB treated**  **mean + SEM** | **Tg-AD DZ/DIB treated**  **mean + SEM** | **p-value** | **F(DFn, DFd)** |
| --- | --- | --- | --- | --- | --- | --- |
| Hippocampus | 2.24E-04 + 4.36E-06 | 2.62E-04 + 1.68E-05 | 2.25E-04 + 8.85E-06 | 2.51E-04 + 1.26E-05 | < 0.01 (g)  0.66 (d) | F (1,34) = 7.67 (g)  F (1,34) = 0.20 (d) |
| SB | 2.07E-04 + 6.63E-06 | 2.42E-04 + 2.00E-05 | 1.96E-04 + 1.33E-05 | 2.25E-04 + 1.33E-05 | 0.03 (g)  0.32 (d) | F (1,34) = 5.09 (g)  F (1,34) = 1.03 (d) |
| CA1 | 2.42E-04 + 4.30E-06 | 2.76E-04 + 1.79E-05 | 2.47E-04 + 4.27E-06 | 2.83E-04 + 1.35E-05 | < 0.01 (g)  0.63 (d) | F (1,34) = 9.21 (g)  F (1,34) = 0.24 (d) |
| CA3 | 2.46E-04 + 6.70E-06 | 2.81E-04 + 1.55E-05 | 2.51E-04 + 5.76E-06 | 2.62E-04 + 8.90E-06 | 0.03 (g)  0.47 (d) | F (1,34) = 5.46 (g)  F (1,34) = 0.53 (d) |
| DG | 2.35E-04 + 5.79E-06 | 2.63E-04 + 2.24E-05 | 2.28E-04 + 4.87E-06 | 2.55E-04 + 1.45E-05 | 0.03 (g)  0.38 (d) | F (1,34) = 5.27 (g)  F (1,34) = 0.80 (d) |

**Supplemental Table 4E:** Reactive Microglia Density (represented as count per square micron) across hippocampal regions at 4 months

**WT untreated	Tg-AD untreated mean**

**Region	mean ± SEM	± SEM	*p*-value	*t*-statistics**

HC	1.15E-05 ± 3.28E-06	1.15E-05 ± 2.29E-06	0.50	t = 0.012, df = 13.47

SB	1.46E-05 ± 4.51E-06	1.67E-06 ± 3.69E-06	0.36	t = 0.359, df = 16.76

CA1	9.33E-06 ± 3.78E-06	8.35E-06 ± 1.79E-06	0.41	t = 0.235, df = 9.99

CA3	9.64E-06 ± 3.54E-06	7.30E-06 ± 1.56E-06	0.28	t = 0.602, df = 9.62

DG	1.16E-05 ± 3.17E-06	1.21E-05 ± 2.91E-06	0.46	t = 0.109, df = 13.90

**Supplemental Table 4F:** Reactive Microglia Density (represented as count per square micron) across hippocampal regions

| **Region** | **WT Untreated mean + SEM** | **Tg-AD untreated mean + SEM** | **WT DZ/DIB treated**  **mean + SEM** | **Tg-AD DZ/DIB treated**  **mean + SEM** | **p-value** | **F(DFn, DFd)** |
| --- | --- | --- | --- | --- | --- | --- |
| Hippocampus | 2.00E-05 + 2.21E-06 | 3.96E-05 + 5.63E-06 | 2.50E-05 + 3.99E-06 | 3.87E-05 + 3.06E-06 | < 0.01 (g)  0.59 (d) | F (1,34) = 18.95 (g)  F (1,34) = 0.30 (d) |
| SB | 2.17E-05 + 2.46E-06 | 4.18E-05 + 3.13E-05 | 2.48E-05 + 4.19E-06 | 4.89E-05 + 4.71E-06 | < 0.01 (g)  0.18 (d) | F (1,34) = 35.10 (g)  F (1,34) = 1.86 (d) |
| CA1 | 1.32E-04 + 2.03E-06 | 3.85E-04 + 4.64E-05 | 1.50E-04 + 1.73E-06 | 3.54E-04 + 3.33E-05 | < 0.01 (g)  0.84 (d) | F (1,34) = 53.78 (g)  F (1,34) = 0.04 (d) |
| CA3 | 1.64E-05 + 2.43E-06 | 2.82E-05 + 3.49E-06 | 1.50E-05 + 2.23E-06 | 2.69E-05 + 1.96E-06 | < 0.01 (g)  0.60 (d) | F (1,34) = 20.9 (g)  F (1,34) = 0.29 (d) |
| DG | 1.66E-05 + 2.50E-06 | 4.52E-05 + 5.12E-06 | 2.46E-05 + 2.47E-06 | 4.21E-05 + 4.03E-06 | < 0.01 (g)  0.51 (d) | F (1,34) = 39.25 (g)  F (1,34) = 0.44 (d) |

**Supplemental Table 4G:** Amoeboid Microglia Density (represented as count per square micron) across hippocampal regions at 4 months

**WT untreated	Tg-AD untreated mean**

**Region	mean ± SEM	± SEM	*p*-value	*t*-statistics**

HC	3.99E-06 ± 1.34E-06	5.35E-06 ± 1.46E-06	0.25	t = 0.689, df = 13.90

SB	4.99E-06 ± 1.80E-06	8.60E-06 ± 2.55E-06	0.13	t = 1.157, df = 12.61

CA1	2.72E-06 ± 1.56E-06	2.58E-06 ± 6.86E-07	0.46	t = 0.087, df = 9.60

CA3	2.88E-06 ± 1.33E-06	2.22E-06 ± 6.51E-07	0.33	t = 0.440, df = 10.15

DG	3.50E-06 ± 1.20E-06	3.87E-06 ± 1.17E-06	0.41	t = 0.222, df = 13.99

**Supplemental Table 4H:** Amoeboid Microglia Density (represented as count per square micron) across hippocampal regions

| **Region** | **WT Untreated mean + SEM** | **Tg-AD untreated mean + SEM** | **WT DZ/DIB treated mean + SEM** | **Tg-AD DZ/DIB treated mean + SEM** | **p-value** | **F(DFn, DFd)** |
| --- | --- | --- | --- | --- | --- | --- |
| Hippocampus | 7.96E-06 + 1.21E-06 | 1.46E-05 + 1.91E-06 | 7.58E-06 + 6.97E-07 | 1.50E-05 + 1.18E-06 | < 0.01 (g)  > 0.99 (d) | F (1,34) = 28.19 (g)  F (1,34) < 0.01 (d) |
| SB | 1.06E-06 + 2.02E-06 | 1.72E-05 + 2.39E-06 | 1.02E-05 + 1.79E-06 | 2.33E-05 + 2.84E-06 | < 0.01 (g)  0.23 (d) | F (1,34) = 18.23 (g)  F (1,34) = 1.51 (d) |
| CA1 | 3.77E-06 + 7.50E-07 | 1.24E-05 + 1.33E-06 | 3.80E-06 + 4.52E-07 | 9.94E-06 + 1.07E-07 | < 0.01 (g)  0.22 (d) | F (1,34) = 59.04 (g)  F (1,34) = 1.56 (d) |
| CA3 | 4.50E-06 + 6.52E-07 | 7.53E-06 + 9.02E-07 | 3.65E-06 + 5.97E-07 | 5.97E-06 + 5.21E-07 | < 0.01 (g)  0.09 (d) | F (1,34) = 15.38 (g)  F (1,34) = 3.13 (d) |
| DG | 5.95E-06 + 1.17E-06 | 1.78E-05 + 2.22E-06 | 7.00E-06 + 5.54E-07 | 1.66E-05 + 2.10E-06 | < 0.01 (g)  0.97 (d) | F (1,34) = 41.01 (g)  F (1,34) < 0.01 (d) |

**Supplemental Table 4: Microglia density across the hippocampus.** Values represent the density of microglia throughout the hippocampus and its subregions. Microglia are first analyzed collectively (tables 2A and 2B), then divided into morphologies based on form factor (Supplemental Figure 2), into ramified microglia (tables 2C and 2D), reactive microglia (tables 2E and 2F), and amoeboid microglia (tables 2G and 2H). All data represented show microglia density as microglia count per micron, with data collected from 4-month wild-type and transgenic untreated rats (tables 2A, 2C, 2E, and 2G), and 11-month rats (wild-type and transgenic untreated versus wild-type and transgenic treated with DZ/DIB, tables 2B, 2D, 2F, and 2H). Data in tables 2A, 2C, 2E, and 2G are analyzed with one-tailed unpaired t-tests with Welch’s corrections, while data in tables 2B, 2D, 2F, and 2H are analyzed by ordinary two-way ANOVA with Sidak’s post-hoc tests. Abbreviations: Tg-AD – transgenic model of Alzheimer’s disease, DZ / DIB – diazoxide / dibenozylmethane; WT – wild-type, g – genotype effect; d = DZ/DIB drug treatment effect, SEM = standard error of the mean, DG – dentate gyrus, CA – cornu ammonis, HC – hippocampus. (g) = genotype; (d) = drug treatment.
